# Development of KASP Molecular Markers and Construction of Fingerprinting for Cowpea(*Vigna unguiculata (L.) Walp*) unguiculata Based on ddRAD-Seq

**DOI:** 10.64898/2026.02.12.705477

**Authors:** Juan Xiang, Zhuoling Zhong, Chengming Zhang, Min He, Kun Cai, Lanping Gu, li Xu, Shilin Su, Yi Zou, Jie Li, Kehao Cui, Huimin Qiu, Bengang Xian, Shaohong Fu, Ling Chen, Xiaowei Liu

**Author notes:** Corresponding anthor. These authors contributed equally to this work (Juan Xiang, Zhuoling Zhong).

## Abstract

Cowpea (*Vigna unguiculata* (L.) Walp.) is a globally important legume crop. However, the scarcity of efficient molecular markers has hindered molecular breeding efforts and the protection of plant breeders’ rights. In this study, we employed double-digest restriction-site associated DNA sequencing (ddRAD-seq) to characterize the genetic diversity of 19 core cowpea germplasm accessions. We generated 791,621 SNPs, from which 13,469 high-quality SNPs were filtered. Population structure and phylogenetic analyses revealed that these accessions cluster into three distinct groups. To facilitate cost-effective and rapid genotyping, we developed a panel of KASP (Kompetitive Allele Specific PCR) markers. Through rigorous screening for polymorphism and stability, we identified six core KASP markers located in exonic regions. These six markers alone were sufficient to discriminate all 19 accessions. Based on these core markers, we constructed a unique DNA fingerprinting profile and assigned specific QR codes for each accession. This study demonstrates that extracting core KASP markers from ddRAD-seq data is a powerful strategy for germplasm identification. The developed fingerprinting system provides a robust, low-cost tool for seed purity testing, variety authentication, and marker-assisted selection in cowpea breeding programs.

## 1. Introduction

Cowpea (*Vigna unguiculata* (L.) Walp.) is an annual legume crop with high contents of proteins and carbohydrates, and it is also rich in a variety of vitamins and minerals, contributing to antioxidant capacity, anti-aging effects, and the maintenance of normal physiological functions [1]. Cowpea is commonly consumed as a vegetable in Asia, whereas in Africa it is mainly used for its grains [2]. Cowpea is a diploid crop with a chromosome number of 2n = 2x = 22 and a genome size of approximately 620 Mb [3]. Continuous artificial selection during domestication and cultivation has driven genetic variation in multiple traits, resulting in diversified cultivar types [4, 5]. Currently, most cowpea cultivars are selfed lines developed through conventional breeding and are highly similar phenotypically; thus, it is difficult to accurately distinguish cultivars by visual inspection, posing major challenges to traditional distinctness, uniformity, and stability (DUS) testing [6]. Therefore, to facilitate rapid cultivar registration while protecting breeders’ intellectual property, new technologies are urgently needed to support cowpea breeding and variety protection.

Molecular marker technologies are reliable tools for germplasm characterization, cultivar identification, and diversity analysis. Single nucleotide polymorphism (SNP) markers, regarded as third-generation molecular markers, are DNA sequence polymorphisms caused by single-nucleotide variation at the genome level. SNPs are among the most abundant DNA variants in plant genomes and are more likely to be associated with biological functions and phenotypes [7]. Because of their high genetic stability, broad distribution, and convenient detection, SNP markers have been widely used in crop diversity analysis, genetic map construction, association mapping, and marker-assisted selection [8, 26]. Traditional SNP discovery requires substantial labor and time, whereas the development of next-generation sequencing (NGS) has greatly reduced the cost of whole-genome sequencing, promoted genotyping approaches, simplified genome complexity, and enabled the generation of large numbers of high-density SNPs as molecular markers [9]. Kompetitive allele-specific PCR (KASP) is a PCR-based fluorescent SNP genotyping technology developed in recent years. By using fluorescently labeled primers and fluorescence resonance energy transfer (FRET) cassettes to discriminate alleles, KASP offers low cost, high throughput, high accuracy, and high polymorphism. KASP has become one of the most widely used SNP genotyping platforms and marker detection methods, showing great potential for crop genotyping, fingerprint construction, and germplasm identification [10-12]

DNA fingerprinting is an important application of molecular marker technology. By testing cultivars using DNA markers and establishing a profile based on genotypes at each marker, DNA fingerprinting can effectively avoid problems of “same name, different cultivar” and “different names, same cultivar” [13, 14]. This approach has been recommended by the International Union for the Protection of New Varieties of Plants (UPOV) [15]. To date, fingerprinting systems based on KASP-derived SNP markers have been established in many crops, including sweetpotato [16], hemp [17], orchids [15], and cauliflower [18]. In cowpea, extensive efforts have been made to develop genomic resources to facilitate marker-assisted breeding. Muchero et al. developed an Illumina 1536-SNP GoldenGate genotyping array and applied it to 741 recombinant inbred lines from six mapping populations. Approximately 90% of the SNPs were technically successful, yielding 1,375 reliable markers, of which 928 were incorporated into a consensus genetic map [19]. Muñoz et al. [20], based on resources developed from the African cultivar IT97K-499-35 and whole-genome sequencing (WGS) data from 36 additional cowpea populations, supported the development of a 51,128-SNP genotyping assay and generated a consensus genetic map containing 37,372 SNPs. In this study, we used ddRAD-seq (double-digest restriction-site associated DNA sequencing) to identify polymorphic SNP in 19 core cowpea inbred lines, developed KASP markers for genotyping, and constructed DNA fingerprints for these 19 accessions, providing references and evidence for marker-assisted breeding and cultivar identification in cowpea.

## 2. Materials and Methods

### 2.1 Plant Materials

The 19 core cowpea inbred lines used in this study were obtained from the Institute of Horticulture, Chengdu Academy of Agriculture and Forestry Sciences. The characteristics of the 19 accessions are shown in Table 1.

**Table 1.**
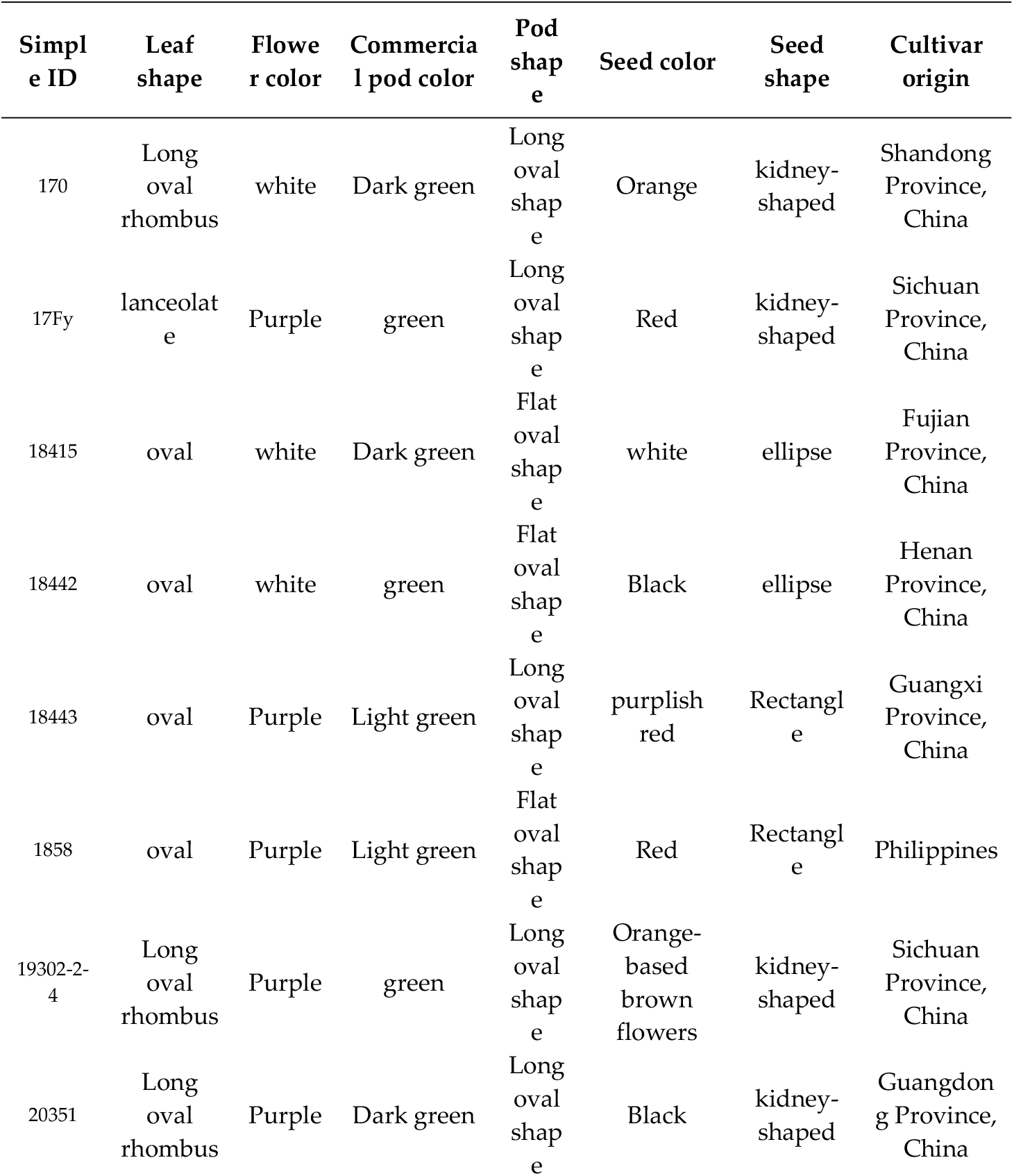
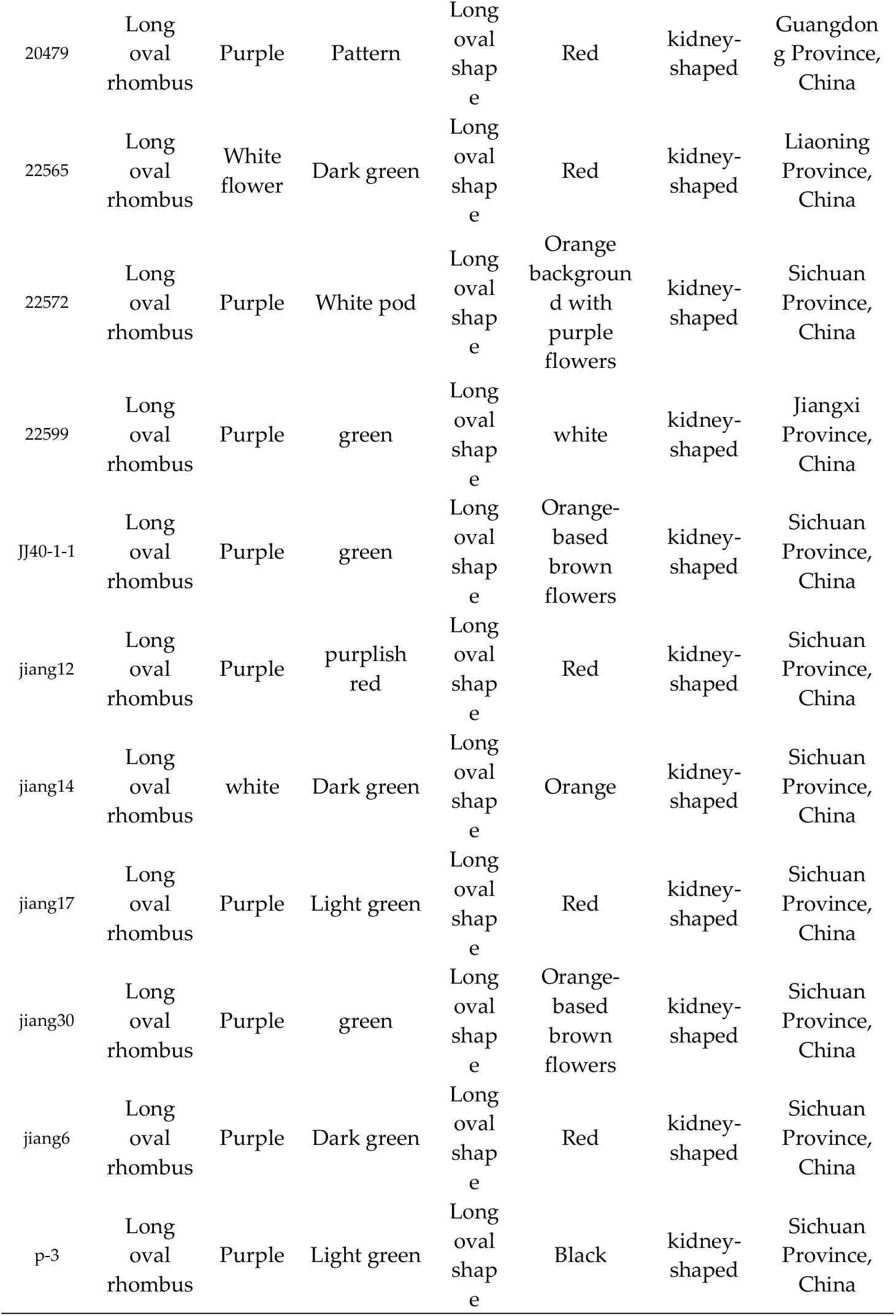
Information of 19 core germplasm resources of cowpea.

| Simpl<br>e ID | Leaf<br>shape | Flowe<br>r color | Commertia<br>l pod color | Pod<br>shap<br>e | Seed color | Seed<br>shape | Cultivar<br>origin |
| --- | --- | --- | --- | --- | --- | --- | --- |
| 170 | Long<br>oval<br>rhombus | white | Dark green | Long<br>oval<br>shap<br>e | Orange | kidney-shaped | Shandong<br>Province,<br>China |
| 17Fy | lanceolat<br>e | Purple | green | Long<br>oval<br>shap<br>e | Red | kidney-shaped | Sichuan<br>Province,<br>China |
| 18415 | oval | white | Dark green | Flat<br>oval<br>shap<br>e | white | ellipse | Fujian<br>Province,<br>China |
| 18442 | oval | white | green | Flat<br>oval<br>shap<br>e | Black | ellipse | Henan<br>Province,<br>China |
| 18443 | oval | Purple | Light green | Long<br>oval<br>shap<br>e | purplish<br>red | Rectangl<br>e | Guangxi<br>Province,<br>China |
| 1858 | oval | Purple | Light green | Flat<br>oval<br>shap<br>e | Red | Rectangl<br>e | Philippines |
| 19302-2-4 | Long<br>oval<br>rhombus | Purple | green | Long<br>oval<br>shap<br>e | Orange-based<br>brown<br>flowers | kidney-shaped | Sichuan<br>Province,<br>China |
| 20351 | Long<br>oval<br>rhombus | Purple | Dark green | Long<br>oval<br>shap<br>e | Black | kidney-shaped | Guangdon<br>g Province,<br>China |
| 20479 | Long oval rhombus | Purple | Pattern | Long oval shape | Red | kidney-shaped | Guangdong Province, China |
| 22565 | Long oval rhombus | White flower | Dark green | Long oval shape | Red | kidney-shaped | Liaoning Province, China |
| 22572 | Long oval rhombus | Purple | White pod | Long oval shape | Orange background with purple flowers | kidney-shaped | Sichuan Province, China |
| 22599 | Long oval rhombus | Purple | green | Long oval shape | white | kidney-shaped | Jiangxi Province, China |
| JJ40-1-1 | Long oval rhombus | Purple | green | Long oval shape | Orange-based brown flowers | kidney-shaped | Sichuan Province, China |
| jiang12 | Long oval rhombus | Purple | purplish red | Long oval shape | Red | kidney-shaped | Sichuan Province, China |
| jiang14 | Long oval rhombus | white | Dark green | Long oval shape | Orange | kidney-shaped | Sichuan Province, China |
| jiang17 | Long oval rhombus | Purple | Light green | Long oval shape | Red | kidney-shaped | Sichuan Province, China |
| jiang30 | Long oval rhombus | Purple | green | Long oval shape | Orange-based brown flowers | kidney-shaped | Sichuan Province, China |
| jiang6 | Long oval rhombus | Purple | Dark green | Long oval shape | Red | kidney-shaped | Sichuan Province, China |
| p-3 | Long oval rhombus | Purple | Light green | Long oval shape | Black | kidney-shaped | Sichuan Province, China |

### 2.2. DNA extraction and re-sequencing

Cowpea were grown in plug trays, when seedlings reached the three-leaf stage, genomic DNA was extracted using a modified CTAB method [21], and DNA quality was assessed by 1.0% agarose gel electrophoresis. DNA concentration was measured using a NanoDrop 2000 UV spectrophotometer, and DNA was then diluted to 50 ng/μL. Double digestion was performed using NlaIII (Hin1II, CATG^) at the Read 1 end and EcoRI (G^AATTC) at the Read 2 end. Qualified DNA samples were used to construct paired-end ddRAD libraries with an insert size range of 300-500 bp. Briefly, 500 ng genomic DNA was incubated with 0.6 U EcoRI (NEB), T4 DNA ligase (NEB), ATP (NEB), and EcoRI adaptors (containing sample-specific index sequences) at 37°C for 3 h, followed by annealing at 65°C for 1 h. Next, NlaIII (NEB) and NlaIII adaptors were added and incubated at 37°C for 3 h. After the reaction, restriction enzymes were inactivated by incubation at 65°C for 30 min in a PCR instrument. Fragment size selection of ligation products was performed by agarose gel electrophoresis, and fragments of 400-600 bp were excised and recovered. Recovered DNA was quantified using Qubit 3.0 (Life Technologies). Twenty-four samples were pooled in equal amounts, and an Illumina TruSeq kit was used to construct the sequencing library. Libraries were sequenced by Genepioneer Biotechnologies(Nanjing, Jiangsu, China).

### 2.3 Raw Sequencing Data Processing and SNP Calling

Perform quality control on the row data to obtain clean data, and then align it to the cowpea reference genome using BWA. The reference genome can be downloaded from https://www.ncbi.nlm.nih.gov/genome/?term=GCA_004118075.2. Utilize GATK to detect Single Nucleotide Polymorphisms (SNPs) and obtain the final set of SNP.

### 2.4 PCA and population stratification analysis

Principal component analysis (PCA) was performed for the identified SNPs using GCTA to infer sample clustering. A phylogenetic tree was constructed using the maximum-likelihood method implemented in FastTree. Population genetic structure was analyzed using ADMIXTURE with the following SNP filtering parameters: mean sequencing depth ≥ 5×, minor allele frequency (MAF) ≥ 0.05, SNP call rate per sample ≥ 0.7, SNP quality score Q ≥ 30, and biallelic loci.

### 2.5 Development of KASP primers

A total of 13,469 SNP markers with no other SNPs within 50 bp upstream or downstream were retained. After applying filters of mean depth ≥ 5×, quality score > 30, minimum completeness > 0.9, minimum minor allele frequency > 0.05, and biallelic status, 11,490 SNP markers were obtained. For each of the 11,490 SNPs, 100 bp upstream and 100 bp downstream sequences were extracted and aligned to the reference genome using BLAST; markers mapping to multiple genomic positions were removed, yielding 6,958 SNP markers. Markers with polymorphism information content (PIC) > 0.35 were retained [22]. Primer3 (v2.4.0) was used to design primers for the 648 SNP markers obtained after the above filtering steps, and loci were further checked to avoid additional polymorphisms before conversion into KASP markers.

### 2.6 SNP Selection, construction of DNA fingerprint and KASP primer validation

Using these 19 cowpea germplasms, the effectiveness of newly developed KASP primers was evaluated. Only loci that generated reliable genotyping results were retained as candidate loci. Filtering based on sequencing results was performed using Perl, and finally six pairs of KASP primers were selected that could discriminate the 19 accessions, enabling construction of a DNA fingerprinting profile based on the core SNP markers. For each primer set, standard FAM and HEX fluorescent tails were added to the 5′ ends of the two allele-specific forward primers. Primers were mixed at a ratio of forward primer 1 : forward primer 2 : common reverse primer = 2 : 2 : 5. Each 20 μL PCR reaction contained 2.8 μL primer mix, 2 μL DNA template, 10 μL master mix, and 5.2 μL ddH2O. PCR was performed in 96-well plates using a touchdown program from 61°C to 55°C: 95°C for 10 min; 10 touchdown cycles of 95°C for 15 s and 61°C for 45 s with the annealing temperature decreasing by 0.6°C per cycle; followed by 28 amplification cycles of 95°C for 15 s and 55°C for 45 s; and finally plate reading at 35°C for 30 s.

## 3. Results

### 3.1 ddRAD-Seq Analysis and SNP Exploration

ddRAD-seq generated a total of 24.51 Gb clean data, after removing low-quality reads and mapping to the reference genome, 170,204,174 reads were retained. The number of reads per sample ranged from 7,938,124 to 12,884,908, with paired-end mapping rates above 87.53%. GC content ranged from 35.24% to 36.17%, Q30 values ranged from 90.64% to 92.91%, and mean coverage depth ranged from 20.07 to 28.41 (Table 2). A total of 791,621 SNP were identified across the 19 cowpea germplasms. The number of homozygous SNPs per accession ranged from 26,948 to 34,834, whereas heterozygous SNPs ranged from 8,641 to 21,439. Among them, jiang17 had the fewest homozygous SNPs, while 1858 had the largest numbers of both homozygous and heterozygous SNPs. The transition/transversion (Ti/Tv) ratio was approximately 2.1 across all accessions (Table 3).

**Table 2.** Statistics on whole-genome sequencing of 19 genotypes and assembly quantity statistics.

| sample | TotalReads | MappedPairedReads | GC (%) | Q30(%) | AverCoverDepth |
| --- | --- | --- | --- | --- | --- |
| 170 | 8519330 | 8368674(98.23%) | 35.72 | 92.81 | 21.11 |
| 17Fy | 9485818 | 8302490(87.53%) | 35.92 | 92.45 | 20.07 |
| 18415 | 8766178 | 8634310(98.50%) | 35.87 | 92.91 | 21.92 |
| 18442 | 8331358 | 8280208(99.39%) | 35.24 | 92.79 | 25.8 |
| 18443 | 8514034 | 8413454(98.82%) | 35.47 | 92.77 | 23.6 |
| 1858 | 8521658 | 8449674(99.16%) | 35.3 | 90.64 | 22.32 |
| 19302-2-4 | 8154588 | 7742220(94.94%) | 36.06 | 92.12 | 22.08 |
| 20351 | 9447530 | 8742348(92.54%) | 35.8 | 92.78 | 23.26 |
| 20479 | 9492924 | 9083562(95.69%) | 36.02 | 92.76 | 23.56 |
| 22565 | 9670652 | 9591158(99.18%) | 35.56 | 92.71 | 25.18 |
| 22572 | 12884908 | 12801680(99.35%) | 35.54 | 92.72 | 28.41 |
| 22599 | 8330818 | 7908320(94.93%) | 35.79 | 92.25 | 22.8 |
| JJ40-1-1 | 8282304 | 8172852(98.68%) | 35.92 | 92.72 | 20.48 |
| jiang12 | 8045084 | 7938958(98.68%) | 36.17 | 92.48 | 21.57 |
| jiang14 | 7938124 | 7703576(97.05%) | 36.06 | 92.62 | 20.31 |
| jiang17 | 8246860 | 8183850(99.24%) | 35.34 | 92.66 | 21.33 |
| jiang30 | 9414600 | 9251762(98.27%) | 35.66 | 92.77 | 22.77 |
| jiang6 | 9579768 | 9430600(98.44%) | 35.7 | 92.84 | 24.69 |
| p-3 | 8577638 | 8515410(99.27%) | 36.05 | 92.76 | 23.29 |

**Table 3.** Statistical table of original variation information.

| sample | SNP number | Transi tion | Transve rsion | Homozygosi ty | Heterozygos ity | Ti/Tv |
| --- | --- | --- | --- | --- | --- | --- |
| 170 | 46072 | 31419 | 14653 | 34443 | 11629 | 2.144202552 |
| 17Fy | 44947 | 30348 | 14599 | 34035 | 10912 | 2.078772519 |
| 18415 | 47771 | 32860 | 14911 | 37651 | 10120 | 2.203742204 |
| 18442 | 36724 | 25082 | 11642 | 27760 | 8964 | 2.154440818 |
| 18443 | 37184 | 25192 | 11992 | 28543 | 8641 | 2.100733823 |
| 1858 | 56273 | 38022 | 18251 | 34834 | 21439 | 2.083283108 |
| 19302-2-4 | 39434 | 26854 | 12580 | 29125 | 10309 | 2.134658188 |
| 20351 | 41892 | 28396 | 13496 | 31548 | 10344 | 2.104030824 |
| 20479 | 44893 | 30637 | 14256 | 33619 | 11274 | 2.149060045 |
| 22565 | 42962 | 29352 | 13610 | 32299 | 10663 | 2.156649522 |
| 22572 | 38394 | 25899 | 12495 | 29234 | 9160 | 2.0727491 |
| 22599 | 37696 | 25560 | 12136 | 27950 | 9746 | 2.106130521 |
| JJ40-1-1 | 41385 | 28122 | 13263 | 30747 | 10638 | 2.120334766 |
| jiang12 | 36942 | 25063 | 11879 | 27349 | 9593 | 2.109857732 |
| jiang14 | 41253 | 28080 | 13173 | 30401 | 10852 | 2.131632885 |
| jiang17 | 36501 | 24683 | 11818 | 26948 | 9553 | 2.088593671 |
| jiang30 | 41526 | 28073 | 13453 | 31048 | 10478 | 2.086746451 |
| jiang6 | 41001 | 28017 | 12984 | 30333 | 10668 | 2.157809612 |
| p-3 | 38771 | 26485 | 12286 | 28433 | 10338 | 2.155705681 |

To further evaluate sequencing quality and the characteristics of SNP, we analyzed sequencing depth distribution, SNP counts and density based on ddRAD-seq data. The average sequencing depth across chromosomes showed an overall stable distribution without obvious depth depletion or excessive enrichment, indicating uniform genome coverage and sufficient depth for SNP discovery (Fig 1A). Significant differences were observed in the number of SNPs among chromosomes: chromosome NC_040289.1 contained the largest number of SNPs (5,588), whereas NC_040280.1 had the fewest SNPs (2,684) (Fig 1B). These differences reflect chromosome-level variation in genetic diversity and provide a basis for prioritizing chromosomes in subsequent marker development. Using a 0.2 Mb sliding window, SNP density analyses revealed heterogeneity among chromosomes. Chromosome NC_040281.1 maintained relatively high SNP density across its length, whereas some chromosomes exhibited local troughs of low SNP density. The color gradient (green to red representing low to high SNP counts) indicated that most windows contained 1-85 SNPs, and only a small number of windows showed highly variable regions with >169 SNPs, suggesting that cowpea genomic variation is broadly distributed but also contains localized hotspots (Fig 1C). Overall, the abundance and clear distribution patterns of SNPs provide a solid molecular foundation for KASP marker development, genetic diversity analyses, and DNA fingerprint construction.

**Fig 1.**
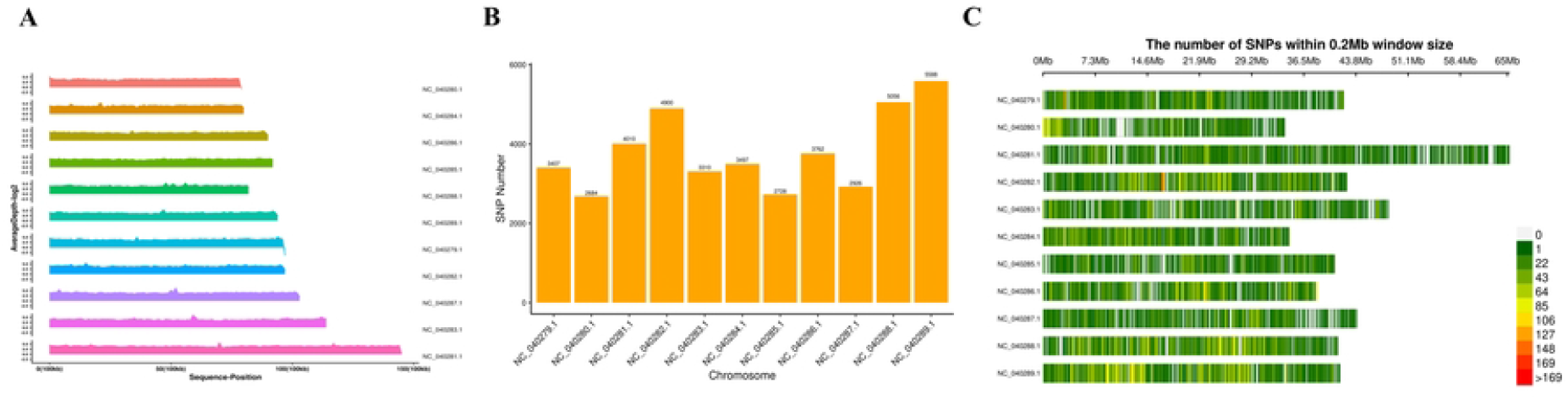
Mutation information statistics. **A**. Depth distribution of ddRAD-seq reads mapped to the cowpea genome. The left y-axis indicates read depth (plotted as log2). The right y-axis shows chromosome names. The x-axis indicates chromosome length. B. Distribution of the number of raw SNP markers across chromosomes. C. Density distribution of raw SNP markers.The x-axis indicates chromosome length, the y-axis indicates chromosome names, and colors represent the number of SNPs within each window.

### 3.2 Genetic Diversity Evaluation of Cowpea Varieties Using SNP Markers from ddRAD-Seq

Principal component analysis (PCA) based on SNP was performed for the 19 cowpea germplasm. The variance explained by principal components PC1, PC2, and PC3 was 31.06%, 22.67%, and 12.62%, respectively, with a cumulative explanation of 66.35% of the total genetic variation, indicating that the first three PCs captured the major genetic differences among accessions. In the two-dimensional scatter plot of PC1 and PC2, the 19 accessions were broadly dispersed without forming clearly separated clusters, suggesting a certain degree of genetic differentiation among accessions. Several accessions showed a wide spread along PC1 but were relatively concentrated along PC2, indicating that PC1 contributed most to distinguishing the accessions. In the three-dimensional distribution including PC3, most accessions showed PC3 values between −0.4 and 0.4, further reflecting differences at specific loci (Fig 2A). Overall, the PCA pattern may be influenced by domestication history, gene flow, and artificial selection, and provides important information for subsequent diversity analyses and core germplasm selection.

**Fig 2.**
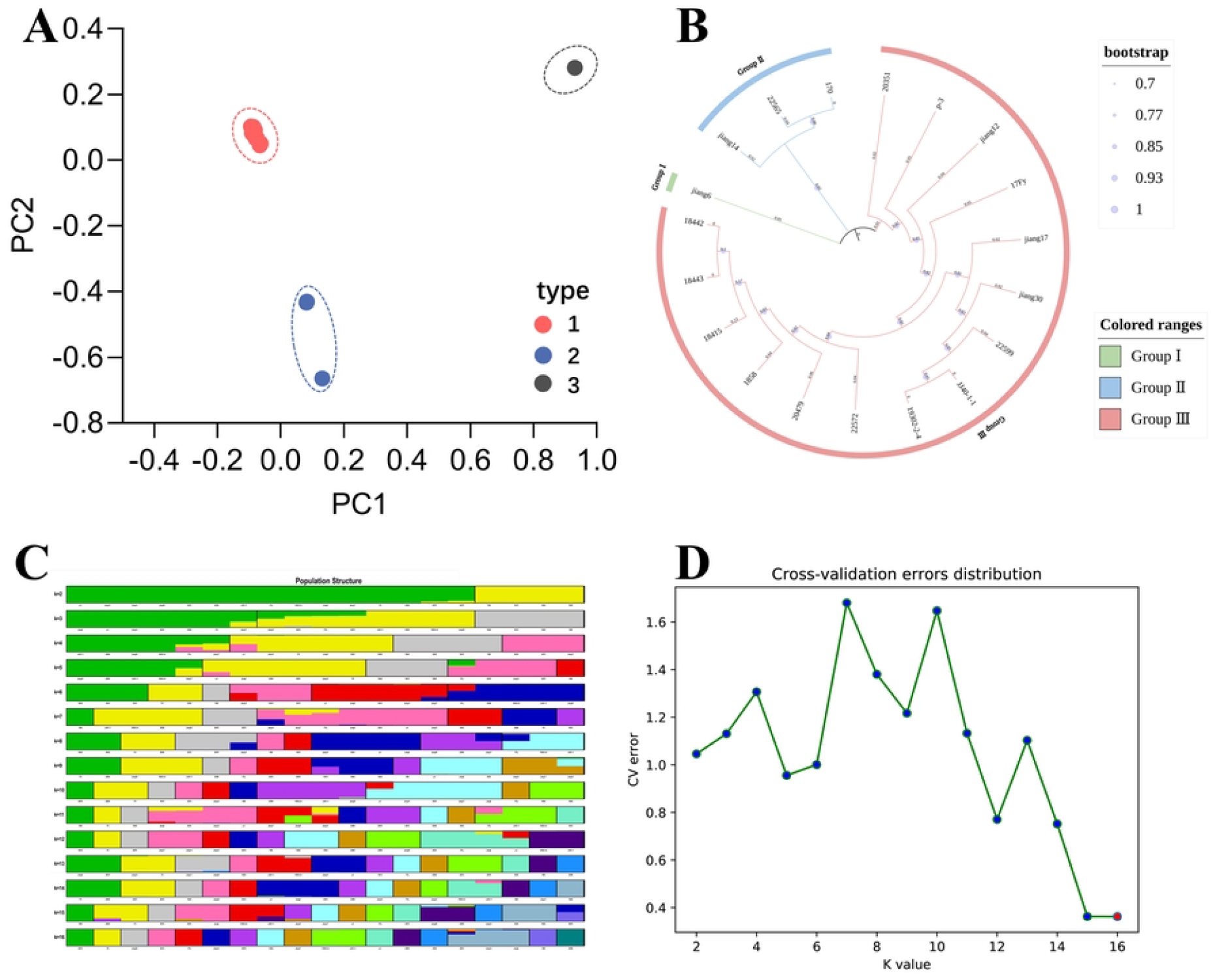
Population genetic analyses of 19 cowpea germplasms based on polymorphic SNP. A, principal component analysis (PCA); B, cross-validation error under different K values; C, population structure of 19 cowpea germplasms at different K values; D, phylogenetic analysis of the population.

Population structure analysis based on ddRAD-seq data indicated that the cross-validation (CV) error reached its minimum when K > 15 (Fig 2B). The inferred genetic structure of the tested cowpea germplasms became progressively more differentiated as the number of assumed ancestral populations (K) increased (Fig 2C). At K = 2, the accessions could be divided into two major genetic components (green and yellow), with most accessions dominated by a single component and only a few showing limited admixture. At K = 3, an additional gray component emerged and the yellow component began to exhibit sub-structure. As K increased from 4 to 16, additional components appeared sequentially (pink, red, blue, purple, etc.), and the genetic backgrounds of the accessions gradually displayed complex mosaic patterns composed of multiple components. Overall, the differentiation was relatively clear at low K values (K = 2-4), reflecting basic genetic groupings, whereas finer differentiation at higher K values (K ≥ 5) revealed potential substructure and gene flow events. These results provide important genetic structure information for defining cowpea genetic groups, mining genetic resources, and constructing breeding populations.

A neighbor-joining (NJ) phylogenetic tree constructed from ddRAD-seq data illustrated the evolutionary relationships and clustering patterns among the 19 cowpea germplasms (Fig 2D). Based on the tree topology, the accessions were divided into three major clades. Group I (green clade) was centered on jiang6 and clustered with 18080, forming a branch that further grouped with Group II. Group II (blue clade) contained jiang14, 22565, 170, and other accessions; this clade showed relatively high bootstrap support, suggesting limited genetic differentiation among these accessions. Group III (red clade) included the largest number of accessions, covering jiang12, 17Fy, jiang17, jiang30, 22599, 18442, 18443, and most other tested accessions. Bootstrap support for pairs such as 18442/18443 and jiang30/22599 reached 100%, and overall support within the clade was high, indicating similar genetic backgrounds and stable clustering. In general, closely related accessions clustered together on the tree, consistent with the diversity results. This further demonstrates the reliability of ddRAD-seq for resolving evolutionary relationships and genetic differentiation in cowpea germplasm, and provides a basis for systematic classification, relatedness identification, and parental selection in molecular breeding.

### 3.3 Design of KASP Primers and Selection of Core SNP Sites

Primer design was performed for the 648 screened SNP markers, resulting in 326 SNP with successful primer design, which were subsequently converted into KASP markers. For these 326 SNP, PIC values ranged from 0.355 to 0.375, and observed heterozygosity ranged from 0 to 1 (S1 Table). These markers were distributed across functional categories including 3′ UTR, 5′ UTR, downstream, exonic, intergenic, intronic, upstream, and upstream/downstream regions (Fig 3A). To evaluate identification efficiency using the 45 exonic-region markers, the identification efficiency increased rapidly from ∼0.20 to 1.00 when the number of markers increased from 0 to 6; when the marker number reached 6 or more, the identification efficiency remained at the saturated level of 1.00 without substantial fluctuation across the full set of 45 markers (Fig 3B). These results indicate that only 6 core KASP markers from exonic regions are sufficient for accurate identification.

**Fig 3.**
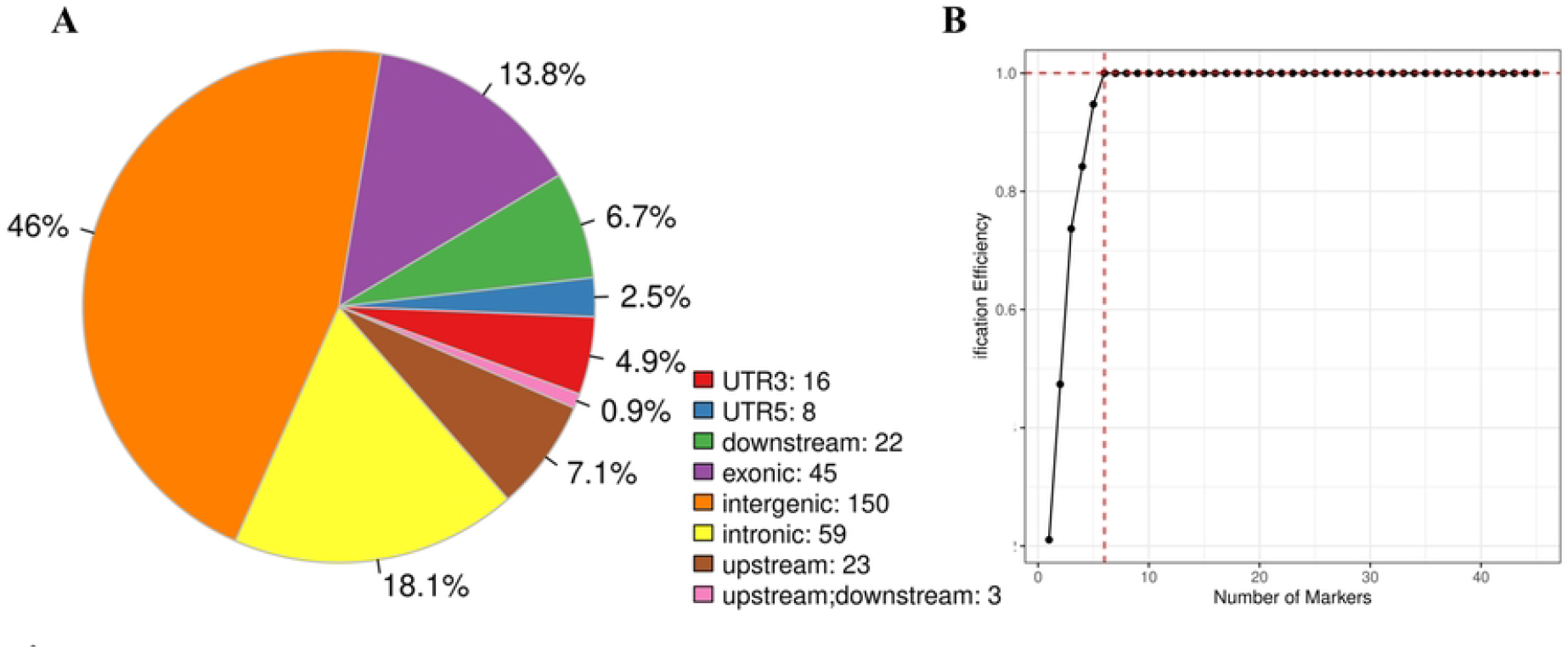
Distribution of SNP markers and marker identification efficiency.

### 3.4 Construction of DNA fingerprints

A fingerprint map constructed based on the six KASP markers (Marker1-Marker6) visually presents the distribution of genotype polymorphisms among the tested cowpea germplasms (Fig 4). The DNA fingerprints of the 19 accessions show that the six KASP markers can be used to discriminate these germplasm resources, effectively distinguishing different accessions. Based on the SNP locus genotype of each variety, each sample was assigned a specific QR code (S1 Fig).

**Fig 4.**
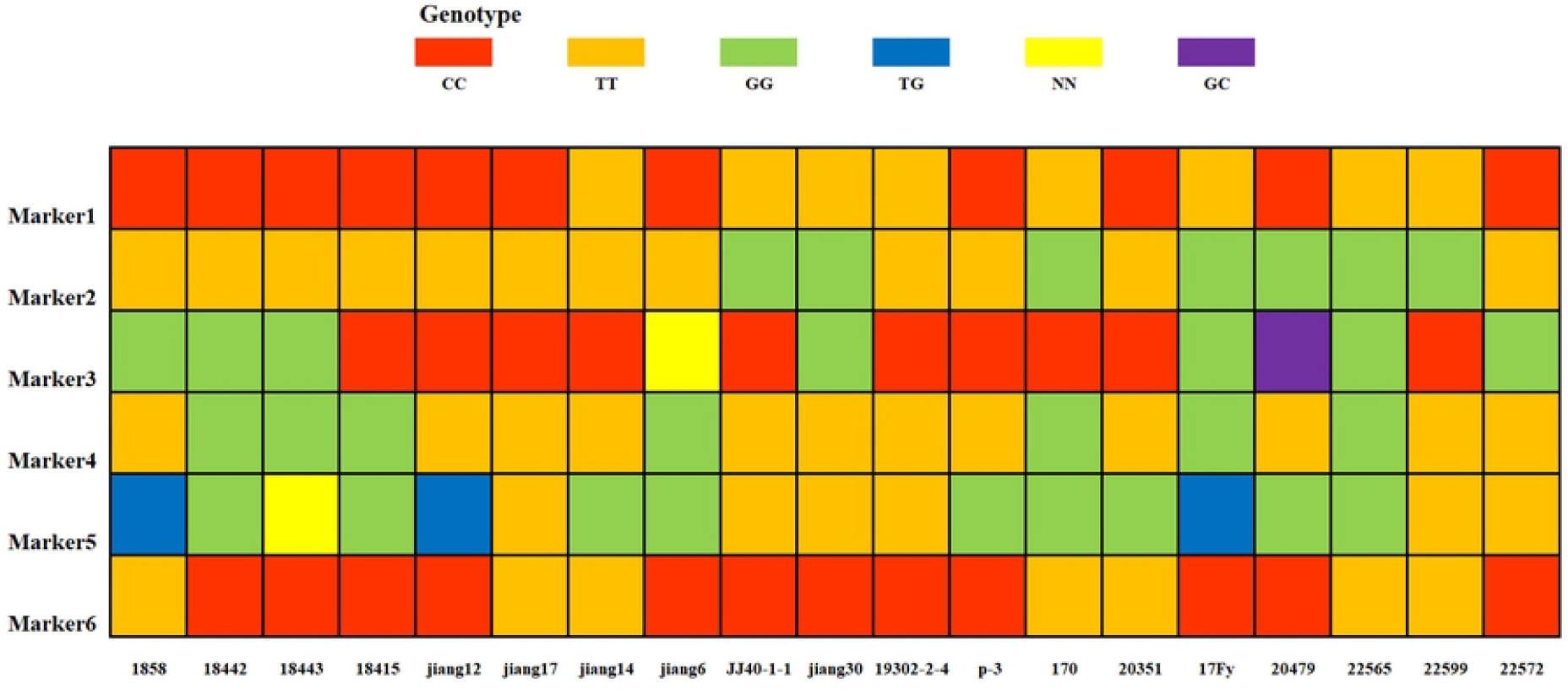
Marker-based fingerprint map. Each row represents one marker and each column represents one sample. Homozygous genotypes (C/C, T/T, G/G, A/) are indicated by red, yellow, and green colors, respectively; heterozygous genotypes are indicated by blue and purple; missing genotypes are indicated by yellow.

From the fingerprint characteristics, the genotypes of each marker differed clearly among accessions. Marker1 and Marker6 were mainly represented by red (homozygous genotype) and yellow (homozygous genotype). Marker2 and Marker4 were mainly green (homozygous genotype) or yellow (homozygous genotype), forming relatively stable genotype groupings across accessions. Marker3 was red (homozygous genotype) or green (homozygous genotype) in most accessions, but purple (heterozygous genotype) and yellow (missing genotype) were also observed, indicating genetic variation and/or missing data for this marker in some accessions. Marker5 also displayed its own distinctive color pattern and discriminative genetic features, with genotype differences observed in some accessions. Overall, the combination of genotypes across the six markers generated unique fingerprint features for the tested accessions, allowing most samples to be distinguished. Only a few accessions (e.g., 18442 and 18443) showed genotype overlap at some markers and may require additional core markers for further discrimination. These results indicate that the selected KASP markers possess good polymorphism and can serve as effective tools for molecular identification of cowpea germplasms, supporting further improvement of fingerprinting systems and accurate germplasm identification.

To further validate the effectiveness of the fingerprinting profile, KASP markers were developed based on the information from six core SNP loci, and genotyping was conducted for 19 cowpea accessions. The results showed that all six markers produced successful genotype calls. All markers were homozygous across the 19 accessions; green and red circles represent two different homozygous genotypes. Marker 1 carried allele C or T; Marker 2 carried allele T or G; Marker 3 carried allele G or C; Markers 4 and 5 carried allele T or G; and Marker 6 carried allele T or C. Overall, based on the genotyping results, these six KASP markers could discriminate the 19 cowpea accessions (Fig 5). This result indicates that the selected KASP markers exhibit good polymorphism and can serve as effective tools for molecular identification of cowpea materials, providing data support for further improvement of the fingerprinting system and accurate identification of germplasm resources.

**Fig 5.**
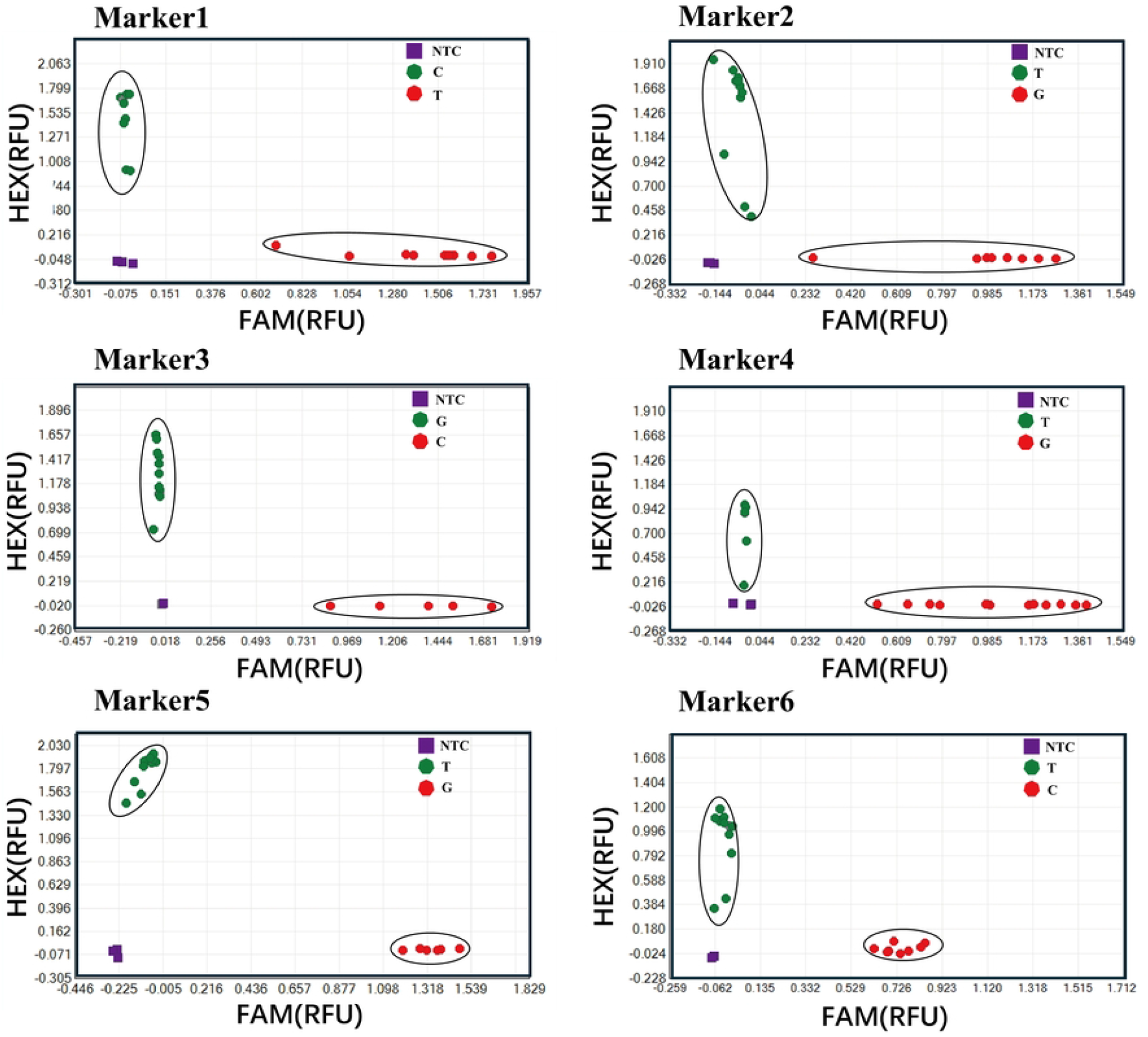
Allelic genotyping of 19 cowpea germplasms using six pairs of KASP primers. Green circles and red circles represent the homozygous genotypes; and the Purple squares on the bottom left of the plot are no template controls.

## 4. Discussion

Cowpea is a globally important legume crop, yet its genetic improvement and variety protection have been constrained by the lack of efficient molecular markers and insufficient population genetic data. As a core reduced-representation sequencing approach, ddRAD-seq is a high-throughput genotyping method based on restriction sites across the whole genome. It sequences DNA fragments generated after complete restriction digestion, and uses two restriction endonucleases to generate fragments of different sizes, thereby increasing the number of genotyped loci [23]. Accordingly, reliable results can be obtained with relatively low sequencing output, reducing potential DNA loss during library construction and effectively decreasing genome complexity, which simplifies library preparation [24]. Therefore, ddRAD-seq has been widely applied in phylogenetics and species identification [25]. In this study, ddRAD-seq was applied to 19 cowpea germplasm accessions, yielding 24.51 Gb of clean data and, after filtering, 170,204,174 reads. Reads per sample ranged from 7,938,124 to 12,884,908; Q30 values were 90.64%–92.91%; GC content was 35.24%–36.17%; and mean sequencing depth was 20.07–28.41. Reads were aligned to the reference genome with a mapping rate exceeding 87.53%. Overall, the sequencing data met high-quality standards and provided a solid basis for subsequent SNP discovery.

Many studies have characterized cowpea germplasm using diverse molecular marker systems; in the present study, we used SNP markers for variety identification. Compared with conventional marker types, SNP markers offer high polymorphism, codominant inheritance, high density, and high throughput, and are widely regarded as ideal DNA markers [26]. Across the 19 core cowpea accessions, a total of 791,621 SNP loci were identified. Their abundance and density exhibited rich polymorphism across the 11 cowpea chromosomes, showing both broadly even distribution and localized hotspots of high variation. For example, chromosome NC_040289.1 contained 5,588 SNPs, whereas NC_040280.1 contained only 2,684 SNPs; this distribution pattern provides a basis for uniformly selecting core markers in subsequent work (Fig 1B). In addition, the transition/transversion (Ti/Tv) ratio was approximately 2.1, consistent with variation patterns in plant genomes and supporting the authenticity of the SNP calls (Table 3).

Developing broadly applicable SNP markers requires substantial background divergence among the tested materials. Based on the SNP loci obtained by sequencing, we performed principal component analysis (PCA) and phylogenetic analysis for the 19 cowpea accessions. The PCA results showed a relatively dispersed distribution of the 19 accessions, indicating a certain degree of genetic differentiation among them (Fig 2A). Phylogenetic analysis further suggested that the 19 accessions could be divided into three branches (Fig 2B). Together, these results indicate appreciable diversity among the accessions, which is favorable for identifying core SNP loci.

Because SNP genotyping often requires complex detection technologies and can be costly, a substantial gap remains between SNP development and practical application in breeding [27]. At present, KASP genotyping based on SNPs has been successfully applied in rice, wheat, Chinese cabbage, and other specie [28-30].KASP substantially reduces genotyping cost while improving practicality, and thus serves as a powerful tool for rapid germplasm identification and marker-assisted breeding [31]. In this study, 648 loci meeting the screening criteria were progressively selected from 791,621 SNP loci, of which 326 were successfully converted into KASP markers, achieving a conversion efficiency of 50.3%. The polymorphism information content (PIC) values ranged from 0.355 to 0.375, indicating strong polymorphism. Functional region annotation showed that the 326 KASP markers covered eight categories, including 3′ UTR, exons, introns, and intergenic regions; exon-located markers accounted for 13.8% (45 markers) (Fig 3A). Such markers are more likely to be associated with phenotypic traits and provide resources for developing functional markers. Marker-number evaluation indicated that when the marker set reached six or more, identification efficiency remained at the saturated level of 1.00 (Fig 3B). After converting six SNP loci into KASP markers, genotyping demonstrated that these six core KASP markers enabled accurate identification of the 19 cowpea accessions and supported the construction of a fingerprinting profile (Fig 4). These results confirm the feasibility and reliability of developing molecular markers using ddRAD-seq reduced-representation sequencing, laying a solid technical foundation for large-scale, high-throughput DNA fingerprinting of additional cowpea varieties and providing technical support for variety authentication, seed genuineness testing, and intellectual property protection in cowpea. Moreover, identifying markers associated with target traits can enable early selection of superior individuals and thereby improve breeding efficiency; the genetic information generated here may inform future germplasm screening and parental combination design. Nevertheless, limitations remain. Only 19 cowpea accessions were included. Although core inbred lines with distinct geographic origins and agronomic traits were selected, the relatively limited sample size may affect the comprehensiveness and representativeness of the genetic diversity analyses. Future studies should expand germplasm sampling to include more accessions with diverse geographic origins and genetic backgrounds to more comprehensively elucidate cowpea genetic diversity and improve the universality and applicability of the fingerprinting system.

## 5. Conclusion

In this study, 19 cowpea germplasms accessions were subjected to high-throughput ddRAD-seq. After strict quality control, 13,469 high-quality SNP were obtained. Through multi-criteria filtering, 326 KASP molecular markers were successfully developed; these markers were evenly distributed across the cowpea genome and exhibited good stability and polymorphism. Six core KASP markers were further selected, and their genotyping results were accurate and reliable. All markers were homozygous in the tested accessions with clear genotype differentiation, enabling effective resolution of genetic backgrounds of cowpea germplasm. Using the 6 core KASP markers, a DNA fingerprinting profile of the 19 accessions was successfully constructed, which can distinguish most tested materials and can be applied in cowpea germplasm identification and marker-assisted breeding. Future work should integrate additional cowpea germplasm resources and develop functional KASP markers tightly linked to key agronomic traits to continuously improve the fingerprinting system and provide more comprehensive technical support for cowpea genetic improvement and sustainable industry development.

## Author Contributions

Juan Xiang, Chengming Zhang conceived and designed the experiments. Min He, Lanping Gu, li Xu, Yi Zou, Kehao Cui, and Huimin Qiu performed the experiments. Shilin Su, Jie Li, Bengang Xian, Shaohong Fu and Ling Chen analyzed the data. Zhuoling Zhong and Kun Cai wrote the manuscript. Xiaowei Liu directed the study and revised the manuscript.

## Informed Consent Statement

All authors have read and agreed to the published version of the manuscript.

## Supporting information

**S1 Fig.**
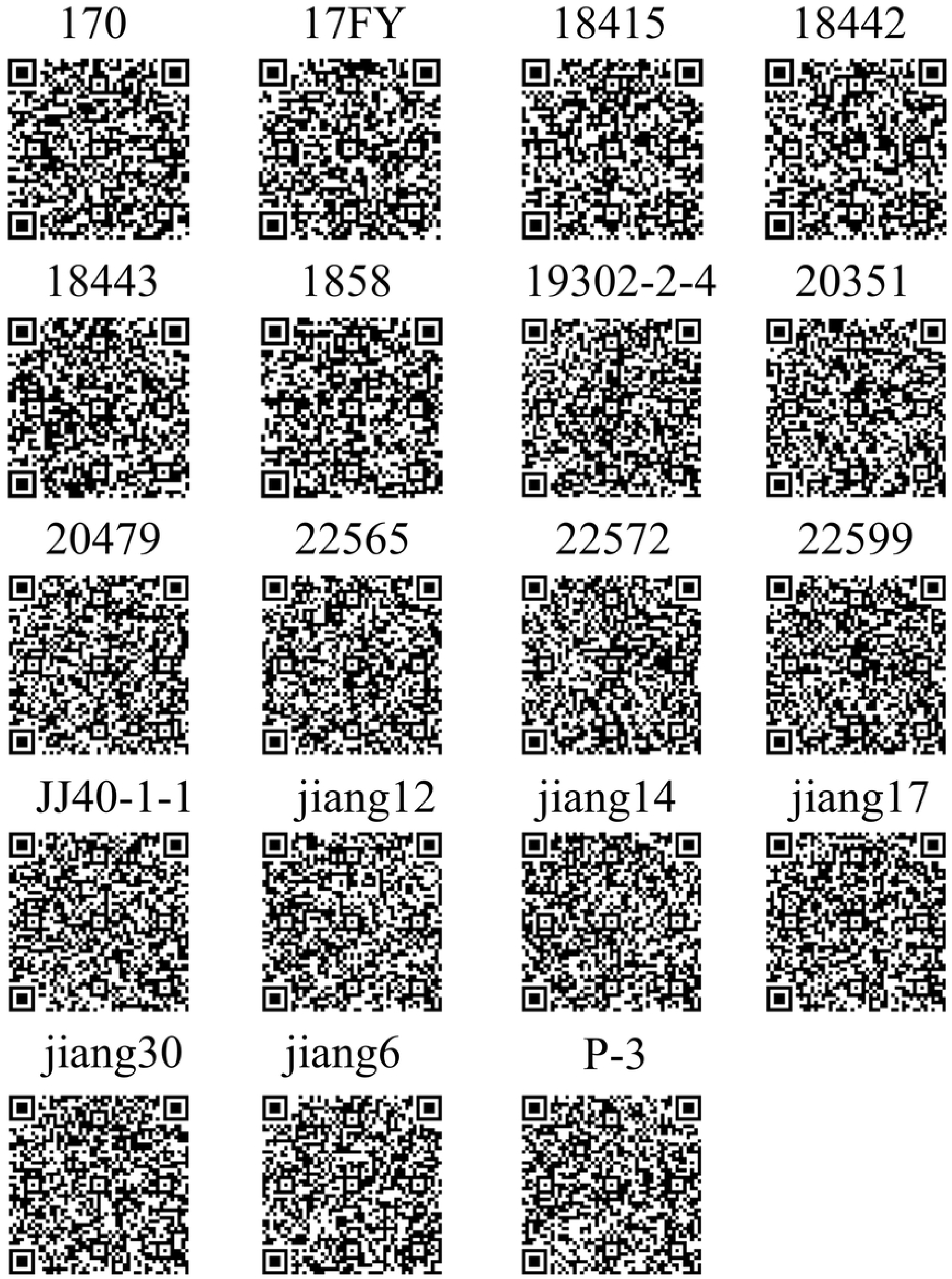
19 accessions of cowpea with two-dimensional barcodes.

**S1 Table.Informations of 326 SNPs**

